# Helix 8 in chemotactic receptors of the complement system

**DOI:** 10.1101/2022.03.07.483401

**Authors:** Szymon Wisniewski, Paulina Dragan, Anna Makal, Dorota Latek

## Abstract

Host response to infection involves activation of the complement system leading to producing of anaphylotoxins C3a and C5a. A complement factor C5a exerts its effect through activation of C5aR1, chemotactic receptor 1, and triggers the G protein-coupled signaling cascade. Orthosteric and allosteric antagonists of C5aR1 are a novel strategy for anti-inflammatory therapies. Here, we discuss recent crystal structures of inactive C5aR1 in terms of an inverted orientation of helix H8, unobserved in other GPCR structures. Analysis of mutual interactions of subunits in the C5aR1 - G protein complex has provided new insights into the activation mechanism of this distinct receptor. By comparison of C5aR1 and its homolog C5aR2 we explained differences between their signaling pathways on the molecular level. A comparison of microsecond MD trajectories started from active and inactive receptor conformations also enabled to elucidate details of local and global changes in the transmembrane domain induced by interactions with the Gα subunit and to explain the impact of inverted H8 on the receptor activation.

## Introduction

Human innate immune system responds to SARS-CoV-2 on various levels, out of which a cytokine storm and an organ damaging, pro-coagulant state, seem to be the most fatal [1,2]. Both these immune system responses are activated by anaphylotoxins C3a and C5a. These effector molecules attract, activate, and regulate innate and adaptive immune cells [3] and lead to formation of membrane attack complexes (MACs). MAC-associated lysis of bacterial membranes is a major role of the complement system composed of more than 30 soluble and surface-expressed proteins [4]. The complement system includes C1-C9 convertases, including C3 and C5 cleaved to C3a and C5a, respectively. Three activation pathways of the complement system: classical (C1, C2, C4), lectin (C2, C4) and alternative (C3) converge at the stage of C3 convertase formation and its cleaving of C3 into C3a and C3b leading to formation of C5 convertase. The final cytolytic MAC is formed by components C5b (from C5), C6, C7 and C8. While MACs disrupt phospholipid bilayers of pathogenic cells, C3b degradation products label cells for phagocytosis, and C3a and C5a anaphylatoxins evoke chemotaxis of immune cells.

An activation of C3aR and C5aR receptors by respective components (C3a and C5a) of the complement system leads to transmitting of proinflammatory signals [4]. Although it is required for the host defense against pathogens, an excessive inflammatory reaction, including overproduction of cytokines, is also a major cause of tissue damaging, e.g., in COVID-19 [5]. Components of the complement system represent druggable targets in treatment of inflammatory diseases, including rheumatoid arthritis and neuroinflammation [6]. The key to success of the complement system therapies is maintaining homeostasis of the immune system rather than turning it off to stop the excessive inflammation [7]. Eculizumab, the anti-C5 monoclonal antibody, is one of few currently used drugs targeting the complement system in treatment of blood-brain-barrier impairments, e.g. neuromyelitis optica. PMX53 and PMX205 are peptide antagonists targeting C5aR1 in treatment of neurological disorders [8]. As shown in phase I and II clinical trials, these two cyclic peptides are well-tolerated and more potent than non-peptide W54011 and JJ47 [9], less violating the Lipinski rule-of-five [10].

Most of C5a effects are mediated through the C5aR1 receptor, while the actual role of C5aR2, the second receptor for C5a which was discovered in 2000, is still under investigation. The cellular localization of C5aR2, either surface or intracellular, is also not clearly defined and highly depends on the cell line/type and experimental conditions [11]. Similarly, discrepancies in C5aR2 ligands, whether it binds only C5a/C5a des Arg or also: C4a, C3a, and their degradation products C4a des Arg and C3a des Arg (ASP), have not been resolved yet. Many studies describe C5aR2 as a dual-acting receptor, which except for its proinflammatory properties also reduces the inflammatory reaction as a decoy receptor trapping the C5a anaphylotoxin and thus limiting the C5aR1 activation [12]. Regulation of the C5aR1 activation [12] also involves carboxypeptidase that cleaves C-terminal Arg and converts C5a into its desarginated form (C5a des Arg). Although C5a des Arg also binds to C5aR1 and C5aR2 [13] it is a less potent anahylotoxin [12] that also triggers other signaling pathways, e.g., HSPC mobilization or lipid metabolism [14].

Functional differences between C5aR1 and C5aR2 have implications in their signaling pathways. C5aR1 transduces the inflammatory signal through the G protein classical pathway while C5aR2 do not interact with G proteins and thus is not considered as a GPCR receptor. However, both receptors efficiently recruit β-arrestins [15] that leads to desensitization. C5aR2 has not been yet characterized but known crystal structures of C5aR1 include helix 8 (H8) of the inverted orientation unobserved in other GPCR structures. Crystal structures of C5aR1 lack the ICL4 loop and C-terminus which could explain whether inverted H8 is an artefact or indeed a distinct feature of chemotactic receptors.

Here, we used crystal structures of C5aR1 [16,17] as templates to generate its inactive conformations. Active conformations of C5aR1 and its complexes with G_i_ protein subunits were generated based on FPR2, the most similar template structure available in PDB. We also performed homology modeling to characterize conformations of its closest homolog C5aR2 and explain its lack of ability to couple G proteins and to affect chemotaxis through the classical G protein-mediated pathway like C5aR1 [15,18]. Based on microsecond MD, we observed that the inverted orientation of amphipathic H8 in C5aR1 did not form any steric hindrance that could prevent interactions of this receptor with G protein subunits. In contrast, H8 in C5aR2 was much less amphipathic and did not form any typical interactions with G protein subunits during simulations, thus confirming its lack of the GPCR-like signaling pathway. In addition, the impact of the G protein subunits only, not involving agonists, on the population of the active C5aR1 receptor conformations were described, following [19].

## Results

### Validation of crystallographic data for C5aR1

An inverted orientation of H8 is present in all three available so far crystal structures of c5aR1 determined independently by two groups: 5O9H [17] and 6C1Q and 6C1R [16]. We performed analysis of these three structures using tools available through PDB and Uppsala EDS server. In all three cases information provided in [16,17] concerning crystallization, structure solution, refinement and validation was consistent with data available in PDB.

In the former case (5O9H), the experimental X-ray data presents over 99% completeness up to the highest resolution shell (2.7Å) with reasonable diffracted intensities (I/σ=7) and satisfactory Rmerge of 19% given that the data set has been combined from 11 partial sets. The above statistics indicates the data as quite reliable. The structural model displays very reasonable discrepancy factor: R-value of 20.8 %, with a very reasonable R-free factor of 23.8%, only slightly higher than R-value. Aside from clashscore and some RSRZ outliers, other structure validation parameters for this deposition exceed the average data quality expected based on PDB statistics relative to both the whole PDB database and the structural subset of similar experimental data resolution. The clashscore is, however, typical for the protein structures of comparable resolution.

The position and conformation of H8, the direction of which is unusual with respect to other GPCRs, is well supported by the electron density distribution. The 2Fo-Fc map at the sigma contour of 1.5 stands in a very good agreement with the atomic positions of the main chain and supports the presence and conformation of side chains such as P287, L289, V293, L294 and in particular T295 and E296 very well (residue numbering according to PDB). The direction of carbonyl groups within the main chain of H8 cannot be doubted. The Fo-Fc residual density map at 1.5 or higher sigma level shows no features in the region of helix-8, indicating a proper model to electron density fit. Electron density of side chains of solvent-exposed R or E residues is less well defined. However, these residues are expected to display considerable disorder.

In the case of 6C1Q and 6C1R [16], the experimental X-ray data presents over 86% completeness up to 2.9Å and over 99% completeness up to 2.2Å with reasonable diffracted intensities (I/σ=6) and very satisfactory Rmerge of 10.6% and 12.8% for 6C1Q and 6C1R accordingly. The latter structure, due to superior completeness and far superior resolution is much more suitable for discussion of the structural details around the helix H8 fragment, although above statistics indicates the datasets for both structures are equally reliable. The structural model for 6C1R displays very reasonable discrepancy factor: R-value of 19.2 %, with a very reasonable R-free factor of 22.4%, only slightly higher than R-value. Aside from RSRZ outliers, other structure validation parameters for this deposition exceed the average data quality expected based on PDB statistics relative to both the whole PDB database and the structural subset of similar experimental data resolution, suggesting a very reliable structural model. RSRZ outliers are not, however, related to the helix H8 fragment.

The position and conformation of H8 is again well supported by the electron density distribution in 6C1R, even more than in the case of 5O9H. The 2Fo-Fc map at the sigma contour of 1.5 stands in very good agreement with the atomic positions of the main chain. In particular, thanks to the better X-ray data resolution, the orientation of the carbonyl groups within the main chain is clearly indicated. The presence and conformation of side chains such as P402, L404, V408, L409 and in particular T410 is also very well supported by the electron density map. The Fo-Fc residual density map at 1.5 or higher sigma level shows no negative features in the region of H8, indicating a proper model to electron density fit. A few positive features indicate that residues N407 and E411 could have been modeled with their side chains (they are replaced by A in the current model, while according to the maps experimental data clearly contains some information about their side chain-positions). Electron density of side chains of solvent-exposed R or E residues is less well defined. However, these residues are again expected to display considerable disorder.

### Differences between C5aR1 and C5aR2 signaling pathways

The described above crystal structures of inactive C5aR1 were lacking a small fragment of ECL2 (an extracellular loop 2), ICL4 (an intracellular loop 4) between TM7 and H8 and C-terminal regions interacting with G protein subunits [15]. These gaps in C5aR1 structures were filled with MODELLER/Rosetta, following procedures described elsewhere [20–22] (see Figure 1A). Then, based on its complete structure, we generated a model of this distinct GPCR receptor in complex with G protein subunits. As a template structure for G protein subunits, we used a crystal structure of a G_i_ protein complex of formyl peptide receptor 2 (FPR2, PDB id: 6OMM) [23] (see Figure 1B). This receptor was the most similar GPCR receptor to C5aR1 that was deposited in PDB in its active conformation interacting with G protein subunits. However, in these homology models of C5aR1 the receptor was still in its inactive conformation, unadjusted to G protein. For this reason, we also prepared [20–22] a second set of homology models that represented an active conformation of C5aR1, based on the FPR2 template (see Figure 1B).

**Figure 1.**
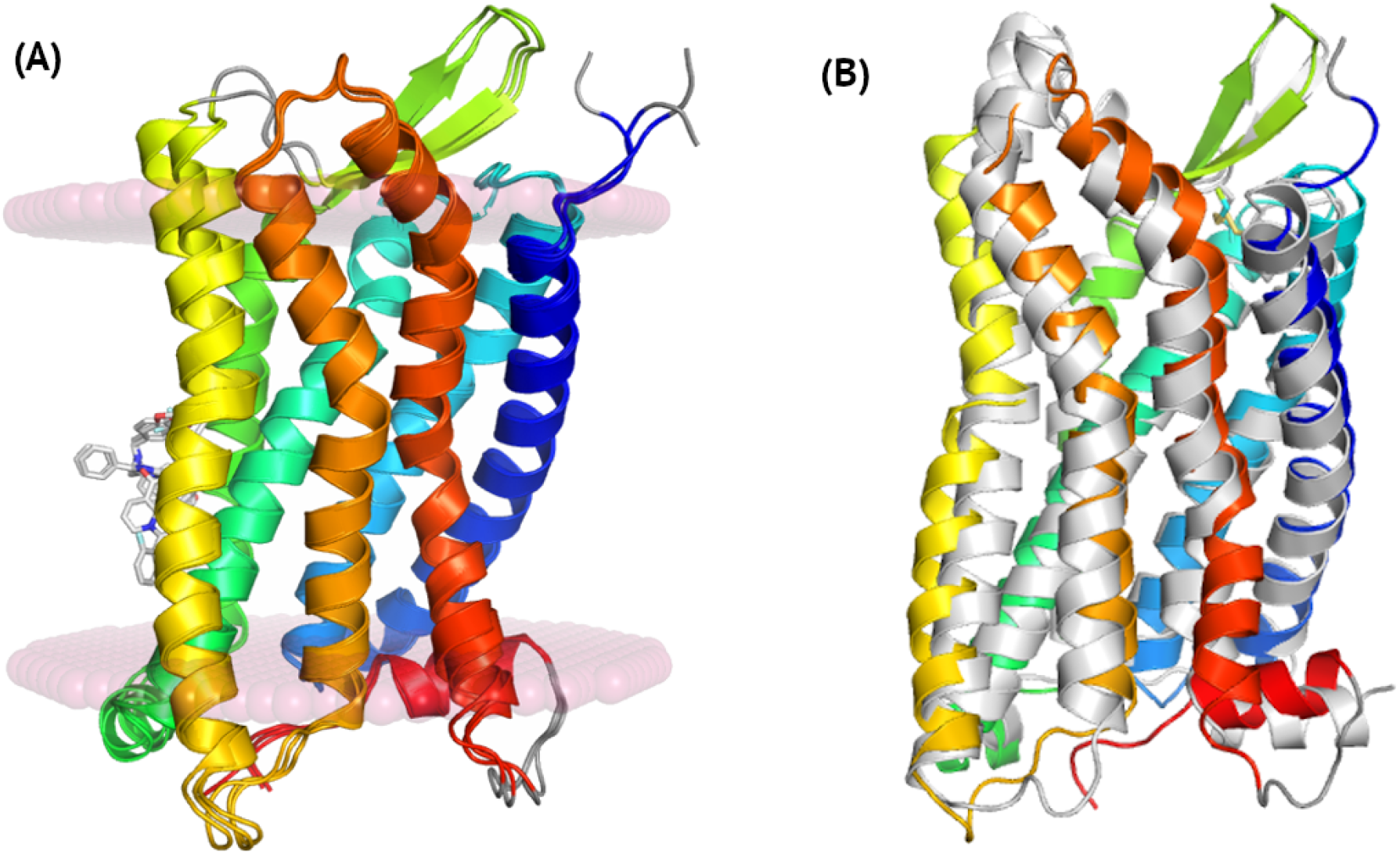
A comparison of inactive conformations of C5aR1 and an active conformation of FPR2. Crystal structures of C5aR1 shown in (A) represent inactive conformations of the receptor, bound to inverse agonists NDT9513727 (5O9H), bound to an orthosteric antagonist PMX53 and to an allosteric antagonist NDT9513727 (6C1Q), and bound to an orthosteric antagonist PMX53 and to an allosteric antagonist avacopan (6C1R). Here, only allosteric antagonists NDT9513727 and avacopan were shown and loops missing in crystal structures were marked with grey. The inactive conformation of C5aR1 (blue-to-red, 5O9H) shown in (B) was compared with the active conformation of FPR2 (grey, 6OMM). An active conformation of C5aR1 based on FPR2 was included in Appendix S1 Figure S1A.

An active conformation of C5aR1 based on FPR2 only slightly changed during microsecond MD simulations (see Appendix S1 Figure S1 and Figure 2). Fluctuations in TMD remained on the level of 2.5 – 3.0 Å with respect to the starting homology model of the complex (see Figure 2C). This level of conformational fluctuations refers to slight changes in loops and side chains of amino acids. Indeed, only a flexible, unstructured ICL4 loop between TM7 and H8 together with EC loops changed their conformations during simulations (see Appendix S1 Figure S1). An intracellular part of TM7 together with ICL4 and H8 refined during first 200 ns of simulations and kept a stable conformation till the end (see Figure 2D). RMSD values of ca. 3 Å were mostly due to changes in ICL4 and slight rotation of the N-terminal part of H8 with respect to the template structure (see Figure 3). Notably, C-terminus of Gα overlapped in both C5aR1 and FPR2 (see Figure 3B). Similarly, H8 in C5aR1 was in the same place, close to Gα, as H8 in FPR2 despite a slight rotation of its N-terminus (see Figure 3B).

**Figure 2.**
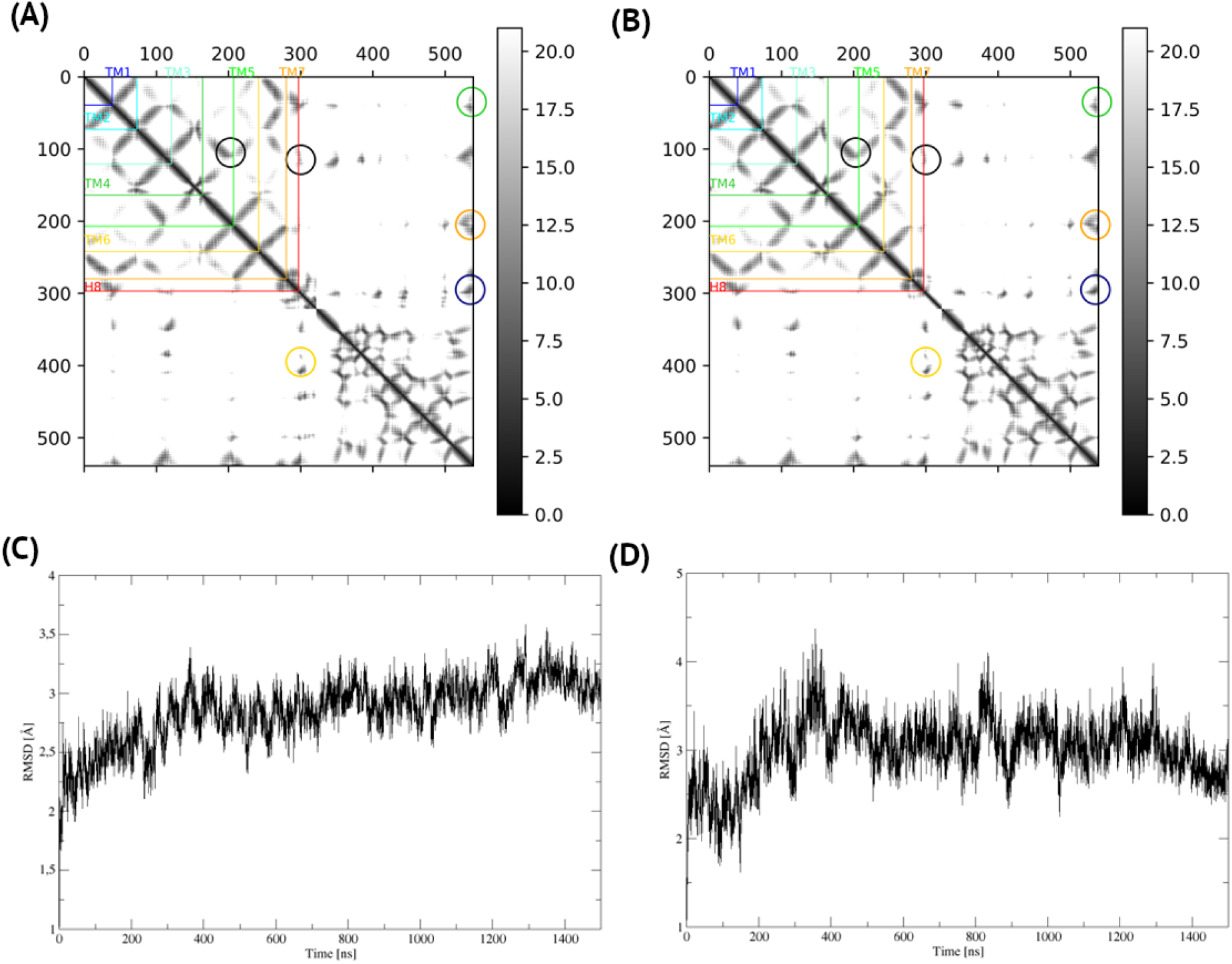
A homology model vs. a microsecond MD-refined model – C5aR1 with G_i_. The homology model of active C5aR1 and G_i_ subunits based on the FPR2 template (A) did not change significantly during microsecond MD simulations (B) which confirmed its reliability. Here, contact maps were shown with the 20 Å cutoff for Cα-Cα distances. The interface shown in Fig. 2 (C5aR1 C-terminus – Gα – Gβ) was not changed significantly (yellow circles). Interactions between C5aR1 and Gα also remained the same (green, orange, dark blue circles), similarly to internal receptor contacts (black circles). The only visible difference was in the receptor region 300-320 (an unstructured, highly flexible C-terminus) that interacted with Gα. (C) The receptor TM core, after quick adjustment to the lipid bilayer during first 20 ns, also remained the same within 3 Å RMSD comparing the starting homology model. The N-terminal part of TM7 and H8 also adjusted during first few ns with the final RMSD not exceeding 3 Å. A gradient contact map for the whole trajectory was included as Appendix S2.

**Figure 3.**
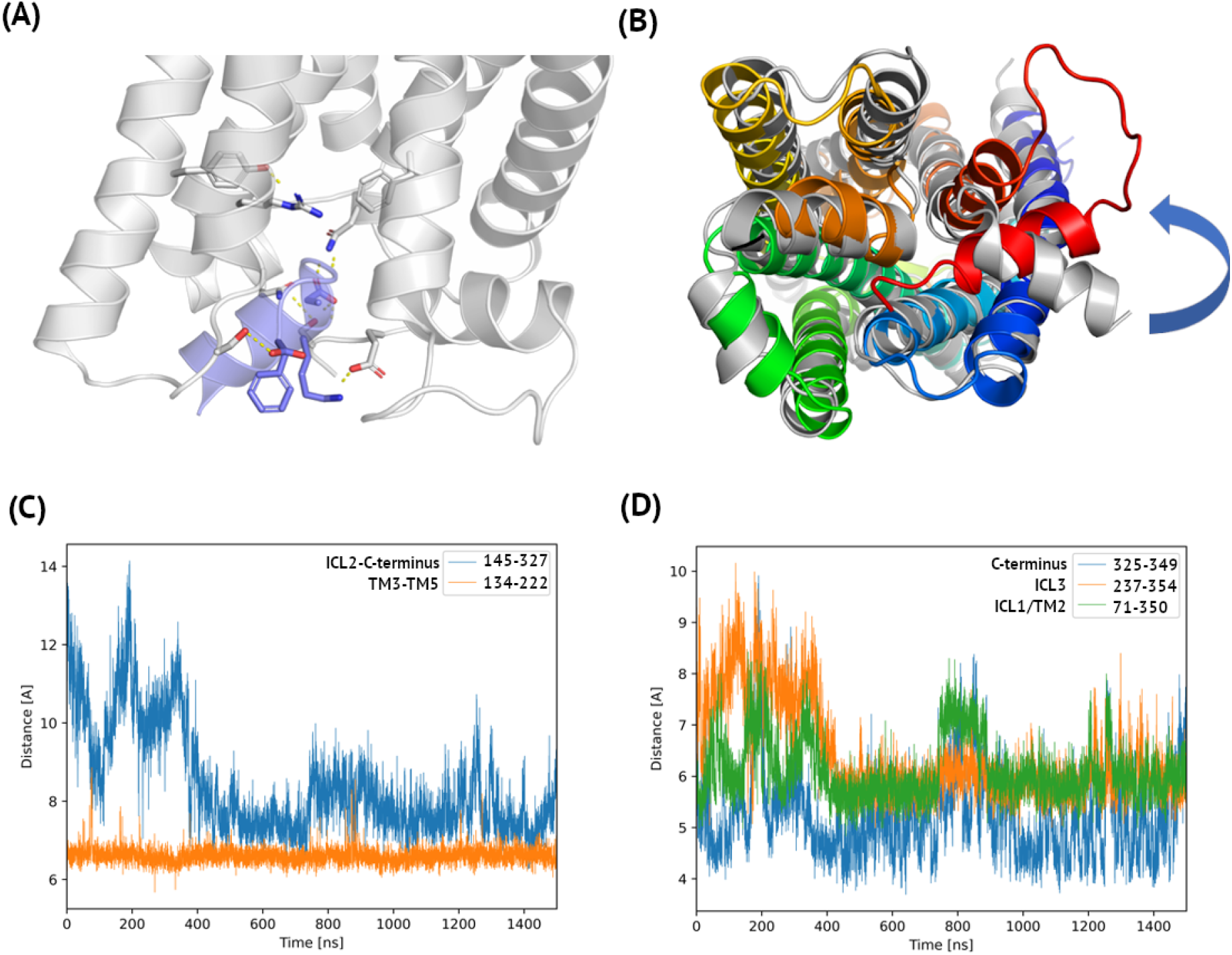
Polar interactions in the complex of C5aR1 and Gα. Here, 1.5 microsecond MD simulations of active C5aR1 was used for analysis of distances between centers of mass of amino acids. In (A) a hydrogen bonding network stabilizing the C5aR1 – Gα interface was shown: Asn71 – Asp350, Glu325 – Lys349, Ser237 – Phe354 (C-cap only) in C5aR1 and Gα, respectively. In addition, intrareceptor interactions were shown: Arg134 – Tyr222 and Gln145 – Ser327. These intrareceptor polar interactions stabilized the TM3 – TM5 interface and the ICL2 – C-terminus region. In (B) the C5aR1 – G_i_ complex refined in microsecond MD was superposed on the FPR2 template (6OMM, an intracellular view). The TM core of C5aR1 was shown in blue-to-red (with C-terminus of Gα in orange) and FPR2 was shown in grey. The C-terminal helix of Gα did not change in comparison with the FPR2 template during simulations. C-terminus of H8 in C5aR1 stayed as close to Gα as H8 in FPR2, but its N-terminus was slightly rotated with respect to FPR2. (C) Distance plots for intrareceptor interactions. After ca. 400 ns of the simulation Ser327 (C-terminus of C5aR1) formed interactions with Gln145 (ICL2). The close distance TM3 – TM5, represented by the distance between residue 134 (the ‘DRF’ motif – equivalent of ‘DRY’) and 222, was observed during the whole simulation. (D) Distance plots for C5aR1 – Gα interactions. Also in this case, after ca. 400 ns Ser237 (ICL3) formed a stable interaction with C-cap of Phe354 (Gα). Interacting Asn71 (TM2) and Asp350 (Gα) additionally stabilized the complex. The least distance, between Lys349 (Gα) and Glu325 (C5aR1 C-terminus), was kept and did not change during the whole 1.5 μs simulation. Here, sequence numbering was adjusted to P21730 and P63096 entries in Uniprot.

Among intramolecular interactions in C5aR1 two hydrogen bonds located in its intracellular part were formed and kept during simulations. These included: Arg134-Tyr222 (TM3-TM5) and Gln145-Ser327 (ICL2-C-terminus) (see Figure 3A and 3C). The former one was observed throughout the simulation while the latter one was formed in ca. 400 ns after the mutual adjustment of the complex subunits and was kept till the end (see Figure 3C). On the interface of C5aR1 and Gα three pairs of amino acids formed stable polar interactions after slight adjustments during first 400 ns. Namely, Asn71 (ICL1/TM2) – Asp350, Glu325 (C-terminus) – Lys349, and Ser237 (ICL3) – Phe354 in C5aR1 and Gα, respectively (see Figure 4C). The latter one involved only the C-cap of Phe354. The former two interactions contributed the most to the stabilization of the complex.

**Figure 4.**
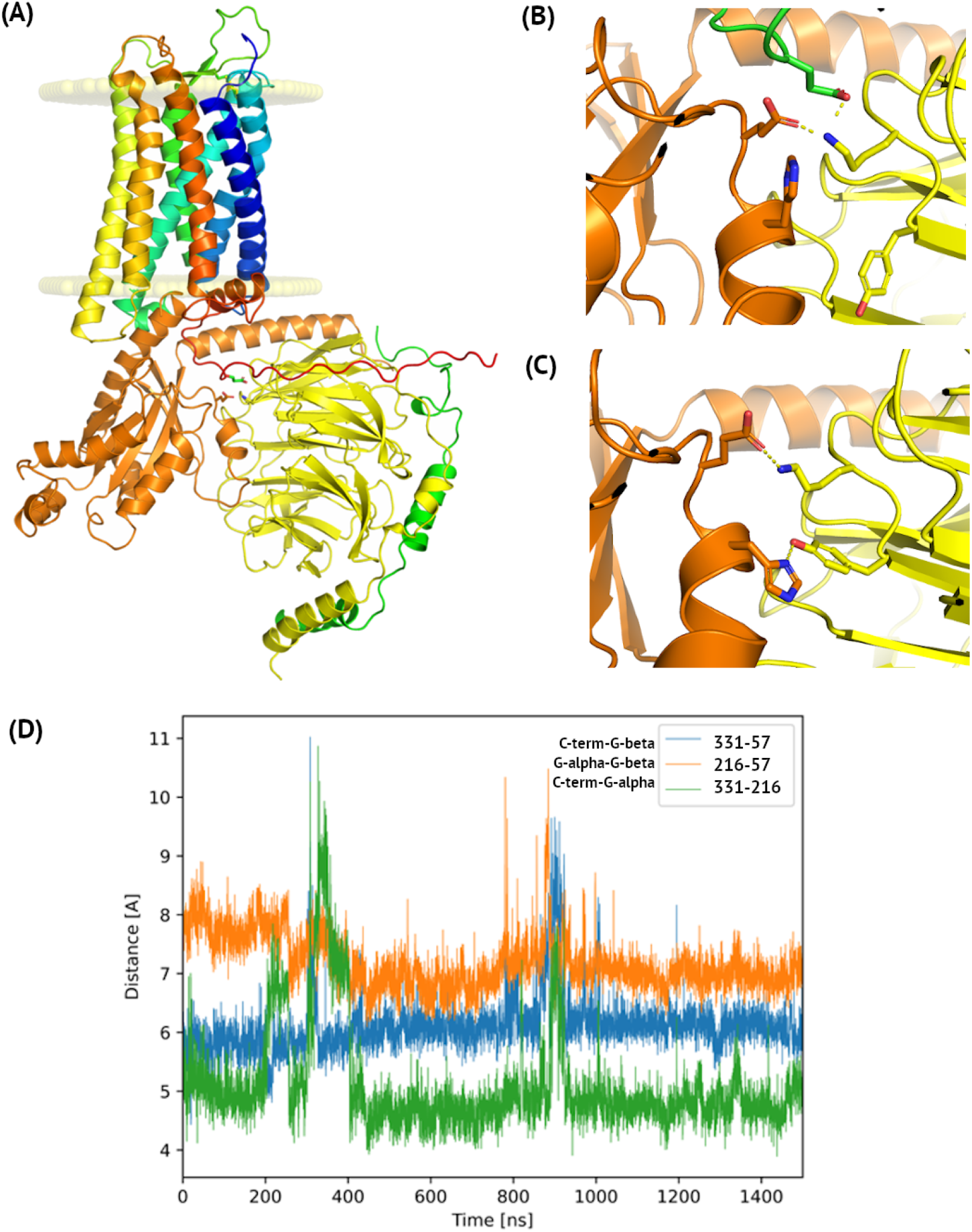
C5aR1 – Gα – Gβ interface observed in microsecond MD simulations. (A) The full simulation system with the indicated lipid bilayer. Here, the structure of the complex was extracted from the last frame of the microsecond MD simulation started from active C5aR1 based on FPR2 with G_i_. A flexible, unstructured C-terminus of C5aR1 (shown in red) did not form any regular secondary structure during the simulation. (B) Details of the C5aR1 – Gα – Gβ interface. Three polar amino acids were close to each other during the whole simulation: Glu331 (green), Glu216 (orange) and Lys57 (yellow) (C5aR1, Gα, and Gβ, respectively). A similar pattern of interface interactions between Glu (Gα) and Lys (Gβ) residues was observed in the cryo-EM structure of active FPR2 with G_i_ (C). Yet, this structure lacks the receptor C-terminus and instead, only an additional hydrogen bond between His (Gα) and Tyr (Gβ) was observed. (B) A distance plot confirming the three-residues’ interaction in C5aR1. Here, sequence numbering was adjusted to P21730 (C5aR1), P63096 (Gα), and P62873 (Gβ) entries in Uniprot.

During the further analysis of the complex subunits in all simulation replicas, we noticed a three-body interaction including the C-terminal region of C5aR1 (see Figure 4) with Lys57 of Gβ in the center. Three amino acids: Glu331, Glu216 and Lys57 in C5aR1, Gα, and Gβ, respectively, were closed to each other throughout the simulation. Among three measured interresidue distances, the Glu331-Glu216 distance was the shortest (4.5-5.0 Å, see Figure 4D). A short distance (ca. 4.5 Å) between centers of mass of Glu331 and Glu216 means that both residues were almost always in a close distance while their negatively charged side chains alternately interacted with positively charged Lys57 (ca. 6 and 7 Å for Glu331 and Glu216, respectively) forming salt bridges. In proteins, two amino acids are in contact if they Cβ atoms are within 8 Å or their Cα within 12 Å [24]. All these distance fullfilled this requirement throughout the simulation and Glu and Lys are know to frequently form salt bridges [25]. Interestingly, three-body interactions were recently described as essential for an accurate prediction of protein folding rates [26]. The same interaction between Glu and Lys amino acids in Gα and Gβ, respectively, was observed in the template structure of FPR2 (see Figure 4C). Yet additionaly, His213 (Gα) interacted with Tyr59 (Gβ). C-terminus of FPR2 was not present in its cryo-EM structure.

The rest of inter-subunits interactions in the C5aR1-G protein complex were kept throughout the simulation which was depicted in Figure 2A-B and Appendix S2 including a gradient contact map for the whole simulation trajectory. Except for the C5aR1 – Gα – Gβ interface described above, also contacts between C5aR1 and Gα were kept (see Figure 2A-B). What is more, no changes were observed in Gα – Gβ and Gβ – Gγ interfaces. The only noticeble conformational change was a slight inwards movement of the N-terminal helix of Gα (see Appendix S1 Figure S2) closer to C-terminus and H8 of C5aR1. However, Gα – Gβ and Gβ – Gγ interactions were kept throughout the simulation (see Figure S2).

Global conformational changes during the receptor activation and subsequent disintegration of the complex and further signal transduction via effector adenylate cyclase are hardly accessible even to microsecond all-atom molecular dynamics simulations. Yet, first steps of the receptor activation can be easily observed in such simulations [27]. Uncovering the driving forces that trigger the signaling cascade provides important clue on how to modify this process with pharmaceuticals. For this reason, we also performed microsecond MD simulations starting from the crystal structure of inactive C5aR1 (see Appendix S1 Figure S3 and Appendix S3 with a gradient contact map). Contact maps for the whole simulation system included similar patterns of interactions before and after simulations (see Appendix S1 Figure S3). However, noticeable changes can be observed in contact maps for TM cores only (see Appendix S1 Figure S3A-B). Contacts between TM6 and other TMs were changed confirming the outward movement of its intracellular part upon the receptor activation. Indeed, the starting, inactive conformation of C5aR1 based on its crystal structure underwent structural rearrangements leading to the semiactive conformation overlapping with the active conformation of C5aR1 based on FPR2 (see Appendix S1 Figure S1).

In a similar way like C5aR1, though never observed [15], we generated an artificial model of C5aR2 – G protein complex to prove its lack of stability and lack of any functional implications. Noteworthy, C-terminus in both C5aR1 and C5aR2 were lacking any regular secondary structure (see the secondary structure prediction described in Methods) and demonstrated a high degree of mobility. This flexibility of C-terminus enables interactions with G proteins and β- arrestin [28]. In a homology model of the C5aR1 – G protein complex (and the artificially generated C5aR2 complex) C-terminus was in a cleft between α and β subunits of G_i_ (see Figure 4A). In the MD-refined complex of C5aR2 its C-terminus was not completely relocated from this cleft but it also did not form the same three-residues interactions observed in C5aR1 (data not shown). TM core and H8 of C5aR2 refined during first 300 ns of simulation and remained stable till the end (see Appendix S1 Figure S4). However, H8 in C5aR2 was moved inside the lipid bilayer during simulations (see Figure 5) because it was less amphipathic than H8 in C5aR1.

**Figure 5.**
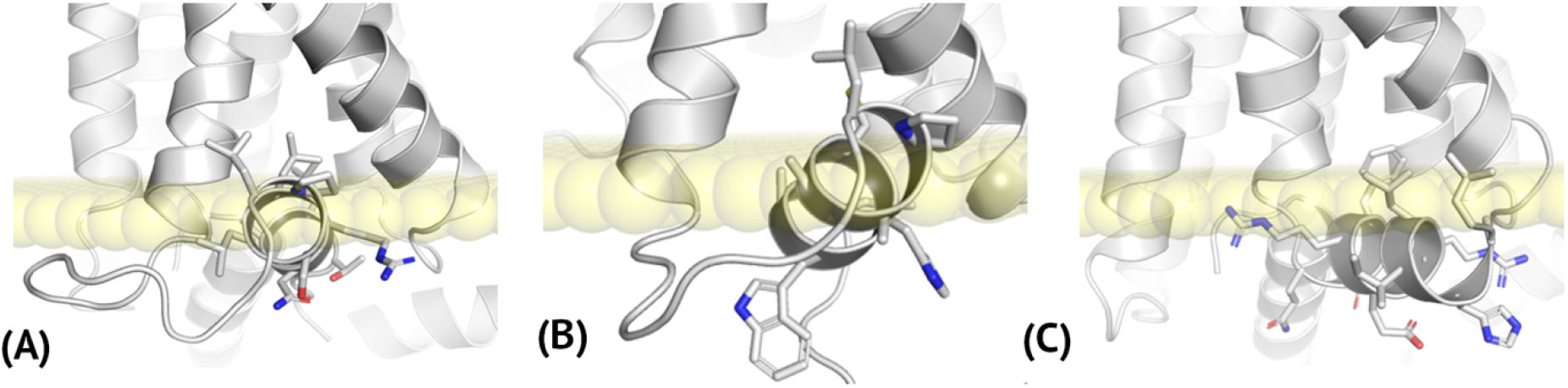
Amino acid composition of H8 in C5aR and FPR2 receptors. (A) C5aR1, (B) C5aR2, (C) FPR1. (A) and (B) structures were extracted from last frames of microsecond MD simulations of C5aR1 and C5aR2, respectively. Here, homology models of active C5aR receptors based on FPR2 were used to prepare simulation systems. In (A) and (C) also interacting Gα subunits were depicted. C5aR2 (B) was not reported to interact with any G proteins. The location of lipid phosphoryl groups was based on inactive C5aR1 deposited in OPM (6C1Q). The length of H8 in all three cases is similar (ca. 10 residues) but the H8 amino acid composition in C5aR1 and FPR2 differs comparing C5aR2. In case of C5aR1 and FPR2 H8 helices are amphipathic like in typical H8 helices of other GPCRs but the content of C5aR2 H8 is mostly hydrophobic on both sites.

The crucial difference between TM cores of C5aR1 and C5aR2 was observed in the region of ICL3 (see Figure 6). C5aR2 includes a shorter loop consisting of 7 residues which implicates also its shorter TM6 and TM5 comparing C5aR1. Also, ICL4 is shorter in C5aR2 by three residues comparing C5aR1. However, both ICL4 loops in C5aR1 and C5aR2 include polar amino acids confirming its outer membrane, intracellular localization (see Appendix S1 Figure S5). Other ECL and ICL loops do not differ to a significant extent (see Figure 6). Yet, Tyr222/His and Arg134/Leu substitutions in C5aR2 disrupt TM3 – TM5 interactions depicted in Figure 3 for C5aR1. Additional disrupting amino acid substitutions in the Gα binding region included: Asn71/Gly and Ser237/Pro (see Figure 6B). Glu325 was not substituted in C5aR2, yet no interactions with the C-terminal helix of Gα were formed during the simulations (see Appendix S1 Figure S6).

**Figure 6.**
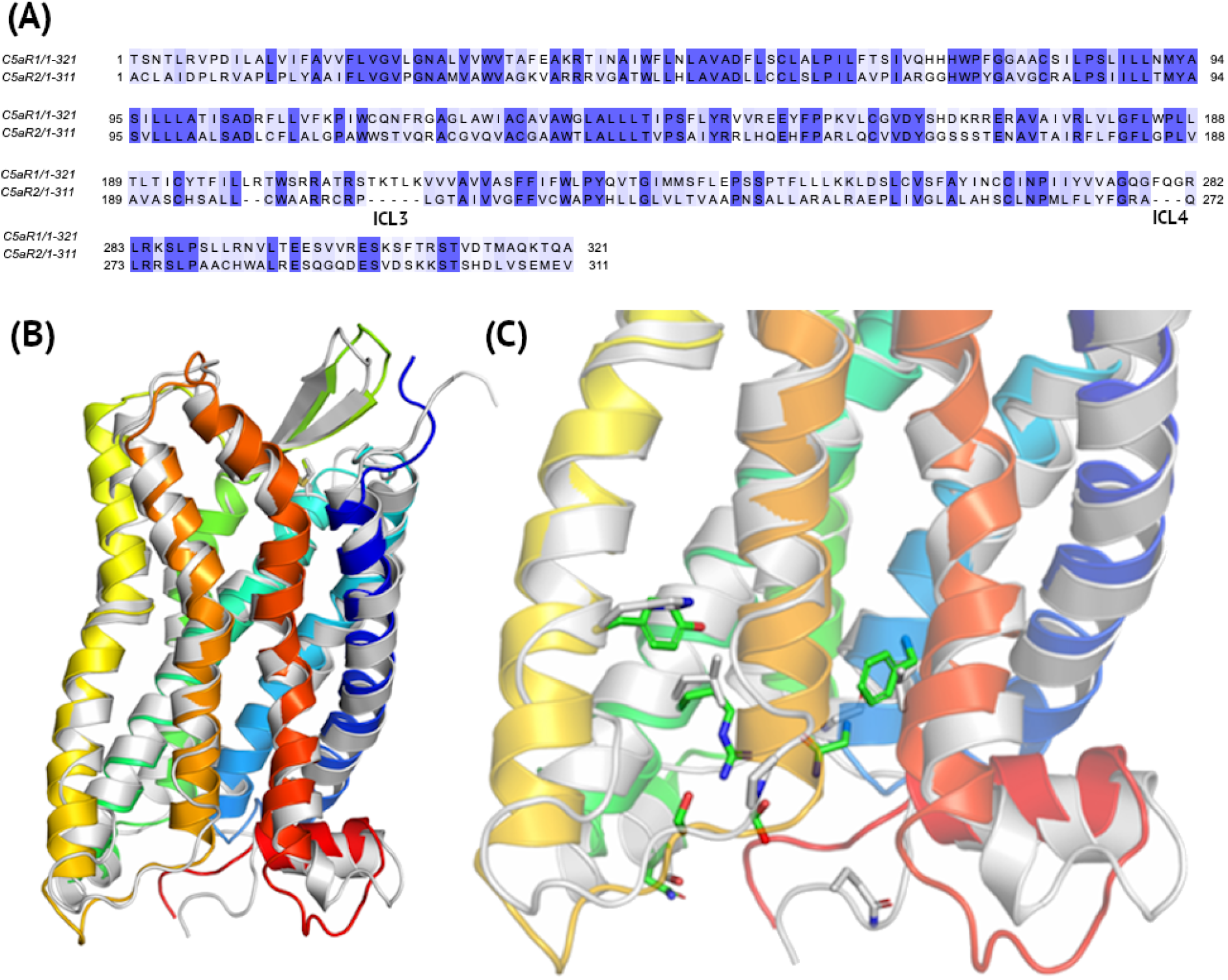
Differences between C5aR1 and C5aR2. (A) – Pairwise sequence alignment with ICL3 and ICL4 loops indicated with 5 and 3-residue deletions in C5aR2. (B) A comparison of the crystal structure of inactive C5aR1 (5O9H, blue-to-red) and a homology model of C5aR2 based on this crystal structure (grey). (D) Amino acid substitutions in the Gα binding region. Here, extracellular 1-29 residues in N-terminus of C5aR1 and 1-27 residues in N-terminus of C5aR2 were truncated relative to sequences deposited in Uniprot (accession numbers P21730 and Q9P296, respectively).

### Helix H8 in C5aR receptors

In either crystal structures of inactive C5aR1 or homology models of inactive C5aR2 there were no steric clashes between H8 and transmembrane helices. Also, in homology models of active C5aR receptors based on FPR2, the inverted orientation of H8 did not overlap with either TMs or Gα. The crystal structure of H8 did not change significantly during MD simulations, regardless the C5aR receptor subtype (see Figure 2D, Appendix S1 Figure S4C-D). This confirms that the inverted orientation of H8 indeed could be a distinct feature of C5aR receptors, not present in other GPCR structures. Nevertheless, amino acid composition of H8 in C5aR1 was more amphipathic-like than in case of C5aR2 (see below) that could also explain differences in signal transduction between these two receptors.

Amphipathic H8 helices in GPCRs are mainly composed from non-polar amino acids interacting with lipids and polar ones facing the cytosol. For example, in H8 of FPR2, Leu and Phe are opposite to His, Ser, Glu, Gln and with two Arg on the border (see Figure 5C). Inverted H8 in C5aR1 is composed from three repetitive Leu residues on the lipids side and Arg, Ser, Asn, Thr on the cytosol side with His and two Leu on the border (see Figure 5A). Yet, H8 in C5aR2 only partially follows this amphipathic amino acid composition. Two Leu and Cys on the non-polar side and Arg and Trp on the polar one with His, Pro and three Ala residues on the border.

Alanine residues inside the helix and only two polar residues on the intracellular side could be the reason why the N-terminal end of H8 in C5aR2 slightly moved into the lipid bilayer side during simulations (see Figure 5B). This could be the reason why interactions between the receptor and Gα were not established (see Appendix S1 Figure S6). What is more, shortening of ICL4 (see Appendix S1 Figure S5) could prevent the proper orientation of H8 with respect to Gα but facilitating interactions of C5aR2 with β-arrestin instead. Noteworthy, in both C5aR1 and C5aR2, ICL4 consists of only polar amino acids (see Appendix S1 Figure S5), confirming the loop-like conformation of this sequence fragment and not amphipathic and helical.

As mentioned above, the amino acid composition of H8 helices in C5aR1, C5aR2, and FPR2 reflected their location relative to the lipid bilayer. Typical, amphipathic H8 in FPR2 was located parallel to the lipid bilayer. A similar location was observed for H8 in C5aR1, although it is slightly more immersed into the membrane comparing H8 in FPR2. In contrast, H8 in C5aR2 is almost completely covered with lipids with only C-terminal residues facing the cytosol (see Figure 5B). What is more, it is not parallel to the lipid bilayer, but its N-terminus, composed of non-polar residues, is moved slightly closer to lipids.

A short helix in ICL4, proposed by the Rosetta cyclic coordinate descent algorithm (see Appendix S1 Figure S7B), unfolded during all simulations. Flexibility of this sequence region was most probably required to properly orient H8 with respect to Gα and TMs (see Figure 3). A helical conformation of ICL4 would stiffen the TM7-ICL4-H8 region and prevent the H8 movement upon the Gα binding during the receptor activation (see below).

### H8 in G protein coupling

As mentioned above, a reliable observation of the GPCR activation requires MD simulations of at least a few microseconds timescale. Yet, our model system remained remarkably stable during all 1.5μs and 1.0μs simulations. Patterns of interactions between C5aR1 and G protein subunits, presented in Figure 2, did not change significantly during microsecond simulations of active C5aR1 conformations. What is more, inactive C5aR1 adjusted its conformation to G protein subunits during simulation (see Appendix S1 Figure S3). TM6 moved away from TM7, and TM3 breaking the inactive state TM3-TM6 lock. C-terminus of C5aR1 and ICL4 adjusted its conformations to fit to the Gα subunit (see Appendix S1 Figure S3A-B). Thus, Gα and other subunits induced conformational changes in the receptor core, as it was suggested in [19]. Namely, Latorraca et al. [19] showed that only after G protein binding the population of active state receptors significantly increases. Agonist binding is not enough to change the distribution of the receptor conformations towards the active state. Nevertheless, much greater simulation timescale is needed to observe further inactive/active state conformational changes. More importantly, simulations starting from an already active conformation of C5aR1 based on FPR2 enabled to observe slight rotation of N-terminus of H8 matching it to Gα (see Figure 3B). This slight rotation of H8 has not been yet observed for simulations starting from inactive C5aR1 (data not shown) as it may require the greater simulation timescale to observe conformations of the fully activated receptor.

Another distinct feature of the C5aR1 – G protein interactions is the three-body interaction (see Figure 4). In the case of C5aR2 this interaction was lost and C-terminus of C5aR2 moved away from these Gα and Gβ residues (see Appendix S1 Figure S6). This loss of interactions with G_i_ subunits was caused by the lack of any stabilizing interactions between TM6 and H8 of C5aR2 and the C-terminal helix in the Gα-1 subunit (see Appendix S1 Figure S6). Shorter ICL3 and ICL4 loops in C5aR2 comparing C5aR1 have impact on the length of TM6 which is much shorter and replaced by too flexible loop (ICL3) which prevents forming stable interactions with Gα (see Figure 6). Also, too short ICL4 prevents the H8 adjustment to Gα in a way that we observed for C5aR1 (see Appendix S1 Figure S6). C-terminus in C5aR receptors is not conserved evolutionary (see Figure 6). A sequence identity on the level of 20 % for less than 30 residues sequences is extremely low. This may additionally explain functional differences between these two receptors. Nevertheless, based on our so far results, only too short ICL3 and ICL4 loops in C5aR2 indeed prevented its coupling to G protein in a way that no GPCR-like signaling pathway could be observed.

## Discussion

C5aR receptors constitute a distinct example of evolutionary diversity of sub-fragments in structures of G protein-coupled receptors. This diversity account not only for intra and extracellular loops but also for the amphipathic helix H8. Many crystal and cryo-EM structures of GPCRs includes disordered H8, e.g, CXCR4 (PDB ID: 3ODU) [29] but none except C5aR1 has included H8 in its inverted orientation so far. Although disordered conformations of H8 could be explained either by experimental conditions or flexibility of this region, a regular helical conformation, 10-residues long cannot be explained only as an experimental artifact.

In this study, we elucidated details of such distinct structural motif and its functional implications. Microsecond MD simulations enabled to observe first steps of the C5aR1 activation and molecular basis of the G protein coupling. Interestingly, these conformational changes were induced only by interactions with G protein subunits, without the presence of the protein agonist C5a, which followed the suggestion of Latorraca et al. [19] on the impact of G protein binding on changes in the inactive/active state populations during the receptor activation. This forms the basis for the next investigations on the full C5a-C5aR1-G protein system involving interactions between C-terminus of C5a and TMs of C5aR1. It would require much larger timescale to observe such propagation of conformational changes of the receptor core. Here, first insights on functional differences between C5aR1 and C5aR2 regarding their signaling pathways were made. Not only global rearrangements in the intracellular part of the receptor were observed, e.g., a rotation of N-terminus of H8 upon Gα binding, but also local conformational changes. Namely, the Glu-Lys-Glu salt bridge in the C5aR1 complex stabilized three-body interactions between C5aR1, Gα and Gβ subunits. Less amphipathic amino acid composition of H8 in C5aR2, disrupting the parallel position to the lipid bilayer, could be one of the reasons why it does not couple with G protein as other GPCRs.

A rational drug design targeting C5aR receptors in treatment of any inflammatory disease requires clarification of their structural and functional differences. Tracking local interactions crucial for the receptor activation and interactions with subsequent components of its signaling pathway is an inevitable step in understanding of these transmembrane proteins of steady interest for pharmacology and medicine. So far, inaccessible by X-ray or cryo-EM regions of H8 and C-terminus of these receptors seems to determine their signaling cascade [30,31] and much more is still to elucidate with constantly released their new structures [32].

## Materials and Methods

### Homology modeling of C5aR receptors

Preliminary models of C5aR1 based on 5O9H, 6C1Q, and 6C1R crystal structures were obtained from GPCRdb [33]. These C5aR1 models were checked for compliance with: Shukla, Robertson et al. 2018, Liu et al 2018, Pandey et al. 2020. Most importantly, it was reported there that C5aR2 is not coupled with G protein but interacts with β-arrestin [15]. Pandey et al. [15] proposed that differences between C5aR1 and C5aR2 regarding the G protein/arrestin signaling pathways were mostly due to this inverted orientation of H8. Previous studies described in [15] suggested that since β-arrestin interacts with both C5aR receptors the loop region between TMH7 and H8 (named ICL4) is rather of irregular structure. N-terminus should not interact with any ligand, while C-terminus is most probably disordered. It was confirmed by additional secondary structure/disorder regions predictions by RaptorX & Robetta [34,35] (see Appendix S1 Figure S9).

GPCRdb models included a typical orientation of H8 in contrast to PDB structures of C5aR1. The inverted helix H8 has been added based on 6C1Q PDB structures because 5O9H included mutated residues in this region. Missing residues in TMH7 were refined based on the 5O9H structure. Crystal structures of C5aR1 end at R330, similarly to GPCRdb models. But residues that are important for G protein or β-arrestin binding include: Ser334-Thr339, Thr336-Thr342 [15–17]. For this reason, C-terminus was added in an unstructured, extended conformation by MODELLER.

Based on these preliminary structures of C5aR1 from GPCRdb which were based on: 5O9H, 6C1Q, and 6C1R we generated full-length models of this receptor using: MODELLER [36], Rosetta [37,38], and GPCRM [20–22]. G_i_ subunits from the template cryo-EM structure of FPR2 (PDB id: 6OMM) were included using MODELLER. In total, 500 TM core models were generated by MODELLER, followed by loop modeling in Rosetta (CCD, 5000 models). 50 top-scoring models were subjected for further analysis. Rosetta models with refined ICL loops that were missing in crystal structures of C5aR1 were subjected to short MD simulations (20-100 ns) to select the most stable conformation of this receptor (data not shown). Two (one helical and the other unstructured) ICL4 loops demonstrated the least RMSD fluctuations and were selected for MD replicas. We selected two versions of ICL4 because of differences in results of loop modeling by Rosetta-CCD [39] and Rosetta-KIC [40] (see Appendix S1 Figure S7). The former algorithm favorized regular, helical conformations of the loop.

During the next step loop models of C5aR1 were inspected in Maestro (Schrodinger, LLC) for protonation/tautomeric states. In principle, during the described above modeling of missing loops, the remaining fragments from crystal structures were changed as little as possible. Yet, for the following residues rotamers were changed with respect to crystal structures of C5aR1: Lys185, Thr229, Ser237, Val328, Val329. For example, Thr229 and Ser237 rotamers was adjusted to fit the 5O9H PDB structure (6C1Q and 6C1R lacked the ICL3 loop). As mentioned above, C5aR1 models were checked for compliance with [15–17] (see Table 1). Residues mentioned in Table 1 were also indicated in Appendix S1 Figure S8, together with residues mentioned in Figures 3 and 4, for comparison. Especially, the ‘DRF’ motif (referring to ‘DRY’ in other class A GPCRs) is included in both, Table 1 and in Figures 3, similarly to residues involved in the H8 formation (see Appendix S1 Figure S8).

**Table 1.** *The most important residues in C5aR1, based on:* [15–17].

| Function | Residues in C5aR1<sup>1</sup> |
| --- | --- |
| Receptor activation | I124 (I3.40),<br>D133-F135 (‘DRF’ motif)<sup>2</sup> (G protein coupling),<br>W213<br>P214 (P5.50), |
| | F251 (F6.44),<br>F254-Y258 ('FWLPY' motif)<sup>3</sup>,<br>N292,<br>N296-Y300 ('NPIIY' motif)<sup>4</sup> (G protein coupling),<br>S334-T339 (β-arrestin binding),<br>T336-T342 (β-arrestin binding) |
| Interactions with ligands | F44, L92, I96, P113, L115, N119, I124, A128, T129, F135, I155, V159, A160, L163, L166, L167, R175, R178, V186, C188, E199, R206, L209, W213, P214, L215, L218, I220, C221, F224, F251, Y258, T261, D282, V286 |
| Interactions with Na<sup>+</sup> | N296-Y300 ('NPIIY' motif) |
| Water flow | N55, D82, S114, N119, S171, R175, Y192, R206, T217, W255, Y258, N292, N296 |
| Involved in the H8 formation | L57, V58, V61, F75, Y300, V301, L315, L319, V322, L323 |

<sup>1</sup>Sequence numbering is the same as in the P21730 entry in Uniprot.
<sup>2</sup> Refers to 'DRY' in other class A GPCRs
<sup>3</sup> Refers to 'CWxPY' in other class A GPCRs
<sup>4</sup> Refers to 'NPxxY' in other class A GPCRs

Complete, curated models of C5aR1 from Maestro were subjected to the HOMOLWAT server [41]. Minor changes to remove steric clashes were made to positions of water molecules proposed by this server. The complete models of inactive and active C5aR1 were also used to generate homology models of C5aR2. Both, inactive and active C5aR2 conformations were generated with MODELLER. As a result of the described above procedure, six replicas in total – four replicas for C5aR1 (two for active and two for inactive receptor conformations, differing mostly in ICL4 loop conformations) and two replicas for C5aR2 (one for inactive and one for active) were used to generate simulation systems.

### Molecular dynamics simulations of C5aR receptors

Complexes of C5aR receptors were prepared for molecular dynamics simulations using the CHARMM-GUI web server [42] including the conserved ECL2 disulphide bond. Each of six simulation systems, containing the receptor complex embedded with the lipid bilayer (OPM-oriented [43]) and solvated (TIP3P), was neutralized by addition of Na^+^ and Cl^-^ ions, with a typical ionic concentration of 0.15 M. The bilayer was formed by POPC and cholesterol molecules with proportion of 3:1. The number of atoms in each simulation was equal to circa 194000 atoms. The Charmm36 force field was used in each simulation, since this force field was tested in various aspects of full-atom simulations of biological systems composed of membrane proteins [44–47]. The equilibration step included six stages, lasting for: 20 ps (steepest descent minimization), 250 ps (conjugated gradients minimization), 250 ps, 250 ps, 500 ps, 500 ps, and 500 ps. During six equilibration stages atomic position restraints were gradually released, e.g., for the protein backbone atoms: from 10 (1st stage) to 0.1 kcal·mol-1·Å-2 (6th stage). The first two stages were performed in NVT, the next four in NPT (1 bar, 303.15 K) using the Langevin dynamics. The production run in NPT was performed using the Langevin piston Nose-Hoover method (1 bar, 303.15 K) and lasted more than 1 μs (or 1.5μs) for each system. The GPU version of NAMD was used for all MD simulations [48], every tenth frame was taken for analysis. TM cores quickly adapted to the lipid bilayer during first 20 ns of production runs. Yet, conformational fluctuations of the whole complex stabilized only after about 300 ns of production runs with the heavy atom backbone RMSD of TMs equal to about 3 Å (C5aR1) 4 Å (C5aR2) and with respect to starting homology models and did not change after further extension of the simulation time (see Figure 2C-D and Appendix S1 Figure S4A-B). Noteworthy, a small helix located in ICL4 as proposed by Rosetta CCD (see Appendix S1 Figure S7B) was not maintained in all MD simulations even though it was highly populated among Rosetta-generated conformations and was assigned the lowest energy due to its regular secondary structure. None of MD replicas includes this helical conformation of ICL4 at the end of the simulation.

## Authors contributions

Conceptualization – DL; data curation – SW, DL; formal analysis – SW, DL; funding acquisition – DL; investigation – SW, AM, DL; methodology – SW, DL; project administration – DL; resources – SW, DL; software – SW, DL; supervision – DL; validation – SW, AM, DL; visualization – SW, DL; writing – original draft preparation – SW, AM, DL; writing – review & editing – SW, AM, PD, DL.

## Acknowledgements

DL acknowledges National Science Centre in Poland (2020/39/B/NZ2/00584) and computational support from the University of Warsaw Biological and Chemical Research Centre. We thank David Aranda Garcia and Jana Selent from: GPCR Drug Discovery Lab, Research Programme on Biomedical Informatics (GRIB), Hospital del Mar Medical Research Institute (IMIM) – Department of Experimental and Health Sciences of Pompeu Fabra University (UPF) for valuable comments.

## Supporting information

**S1 Appendix. Additional figures.** Figures that were not included in the main text.

**S1 Figure. Global changes of C5aR1 induced by G_i_ subunits.**

**S2 Figure. A homology model vs. a microsecond MD-refined model of active C5aR1 – the location of G protein subunits.**

**S3 Figure. First steps of the C5aR1 activation observed in microsecond MD simulations.**

**S4 Figure. Results of microsecond MD simulations performed for C5aR2.**

**S5 Figure. Amino acid composition of ICL4 loops in C5aR receptors.**

**S6 Figure. Loss of crucial interactions between C5aR2 and G_i_ subunits.**

**S7 Figure. A comparison of loop modeling algorithms in Rosetta.**

**S8 Figure. Residues important for C5aR1 function.**

**S9 Figure. Prediction of secondary structure of C5aR1 and C5aR2.**

**S2 Appendix. A gradient contact map for active C5aR1.** Distances were computed between Cα atoms of the receptor and the Gα subunit from each frame of the 1.5 μ MD simulation started from the active conformation of C5aR1 based on FPR2.

**S3 Appendix. A gradient contact map for inactive C5aR1.** Distances were computed between Cα atoms of the receptor from each frame of the 1.5 μ MD simulation started from the cryo-EM, inactive conformation of C5aR1.

## Data availability statement

All data referring to the manuscript, including homology models, MD trajectories, scripts for trajectories analysis are provided upon request from Authors.

